# Can overlooking ‘invisible landscapes’ bias habitat selection estimation and population distribution projections?

**DOI:** 10.1101/2024.01.20.576416

**Authors:** Romain Dejeante, Rémi Lemaire-Patin, Simon Chamaillé-Jammes

**Affiliations:** CEFE, Univ Montpellier, CNRS, EPHE, IRD, Montpellier, France; ISPA, Bordeaux Sciences Agro, INRAE, F-33140, Villenave d’Ornon, France; Mammal Research Institute, Department of Zoology and Entomology, University of Pretoria, Pretoria, South Africa

**Keywords:** species distribution, memory, space familiarity, invisible landscapes, habitat selection, space use, SDM, RSF

## Abstract

Species’ future distributions are commonly predicted using models that link the likelihood of occurrence of individuals to the environment. Although animals’ movements are influenced by physical landscapes and individual experiences (for example space familiarity), species distribution models developed from observations of unknown individuals cannot integrate these latter variables, turning them into ‘invisible landscapes’. In this theoretical study, we address how overlooking ‘invisible landscapes’ impacts the estimation of habitat selection and thereby the projection of future distributions. Overlooking the attraction towards some ‘invisible’ variable consistently led to over-estimating the strength of habitat selection. Consequently, projections of future population distributions were also biased, with animals tracking habitat changes less than predicted. Our results reveal an overlooked challenge faced by correlative species distribution models based on the observation of unknown individuals, whose past experience of the environment is by definition not known. Mechanistic distribution modelling integrating cognitive processes underlying movement should be developed.

## INTRODUCTION

Moving from descriptive to predictive ecology has been a key endeavour for ecologists for decades, especially as the demand for applied science increased. For instance, understanding and predicting the distribution of animals, especially in changing environments, is considered critical to develop appropriate conservation and management practices (Sofaer *et al*. 2019). However, as highlighted by (Mouquet *et al*. 2015), predictions can ‘grow faster than our understanding of ecological systems’, and there is always a great need to try to reduce structural uncertainty of predictive models, among the other sources of uncertainty, as much as possible (Elith *et al*. 2002).

Species distribution models (SDMs) are now commonly used to predict species’ habitat suitability by comparing the environmental characteristics of locations where the individuals are observed and the locations where the individuals could have been. Such models can be developed from snapshot observation data of unknown individuals, in which case the models are generally termed SDMs based on presence/pseudo-absence data (Elith & Leathwick 2009) and estimate ‘habitat suitability’, or from animal tracking data, in which case the models are generally termed resource selection functions (RSF) on used/available data (Boyce *et al*. 1999) and estimate ‘habitat selection’. The difference in terminology hides that the class of models used on the two different types of data are generally similar (Matthiopoulos *et al*. 2015) and fundamentally estimate the disproportionate presence of the species in some habitats.

Habitats are generally defined as a part of the environmental space that is rather homogeneous across a set of variables that are assumed to describe the many relevant aspects of the landscape in which the individuals live. For instance, a habitat will be characterized by having a specific vegetation type, specific slope, and to be at a specific distance to some distinct features of the landscape (e.g. roads).

However, an animal’s selection of where to spend time is not solely influenced by characteristics of locations across the set of environmental layers that are easily observable or measurable by a human analyst, but also by variables as diverse as the landscape of fear (Laundre *et al*. 2010), the social landscape (Webber & Vander Wal 2018; Armansin *et al*. 2020), or the memory landscape (Smouse *et al*. 2010). Although the importance of these landscapes on animal movement is increasingly recognized by ecologists, (especially regarding the influence of predator-prey interactions and spatial memory), their estimation requires long-term tracking of animals (for example to detect which locations are familiar for each individual) and intensive tracking of many individuals (for example to detect the encounters between preys and predators). Therefore, species distribution predicted from RSFs based on tracking data rarely account for these processes, commonly pooling tracked animals together and ignoring individual-level information (Chambault *et al*. 2021; McCabe *et al*. 2021). Similarly, SDMs developed from snapshot observations and species-level occurrence records will necessarily fail to integrate the influence of these processes that need individual-level information to be estimated. Because of the difficulty for researchers to measure and account for these landscapes, which are relevant for animals but do not map directly with physical features of the environment, we termed them ‘invisible landscapes’.

One might however expect that overlooking the influence of these invisible landscapes on animals’ movement decisions may severally impair the accuracy of predictive distribution models. Indeed, describing exhaustively the variables susceptible to influence animal movements is critical to increase the performance of habitat suitability models (Zeller *et al*. 2017). Also, the lack of one variable in the habitat suitability model may bias the estimation of the effect of others (Van Moorter *et al*. 2013). Although many ecologists are aware of the importance of non-physical landscapes (e.g. competition, social interactions, spatial memory…), most predictions of species distributions fail to integrate their influence and there is great need to understand whether systematic biases in predictions are to be expected.

Here we contribute a theoretical study to highlight how accounting, or not, for ‘invisible landscapes’ affects species distribution models. We used an individual-based model to simulate the movement and distribution of animals whose habitat selection was known and influenced by familiarity (i.e. whether a location had been used before by the individual) and recent use by conspecifics. We then investigated to what extent (1) the estimation of the strength of selection for some habitat variables and (2) the prediction of population distribution could be biased when the influence of familiarity and recent use were not accounted for in the analysis. Because SDMs are often used to predict species distribution in changing environments or in future conditions, we used our theoretical approach to test how ‘invisible landscapes’ could bias the prediction of population distribution in such situation. Our results reveal strong consistent biases produced by failing to account for ‘invisible landscapes’, and therefore question the validity of the predictions from SDMs based on snapshot observations of species’ presence, or from RSFs based on GPS data that did not account for individual’s experience, for instance.

## 2. MATERIAL & METHODS

### 2.1. Testing the effect of familiarity and recent use by conspecifics on habitat selection estimation

To test the effect of not accounting for familiarity and recent use of the landscape by conspecifics when estimating habitat selection, we (1) simulated the movements of animals with known selection coefficients for habitat type, familiarity and level of recent use, and (2) compared the ‘true’ coefficient of habitat selection with the one estimated from models that did not contain familiarity and recent use levels as predictors.

#### 2.1.1. Description of landscape, movement model and familiarity and recent use metrics

We developed a model that, for each run, simulates the movements of 500 individuals over a 100×100 habitat raster map during 1000 time steps. The movement of each animal was itself generated using the local Gibbs (LG) movement model proposed by (Michelot *et al*. 2019). The LG model is a step selection model that ensures that the utilization distribution of the animal is consistent with a predefined RSF whose coefficients are used to parametrized the LG model. For each time *t*, a random location was sampled within a 5-cell radius around the current location, 100 potential locations were then uniformly generated within the 5-cell radius around this random location, and the location at time *t+1* was sampled among them with probabilities proportional to the RSF score. In this work, the RSF used to simulate movement of individuals was as follow:

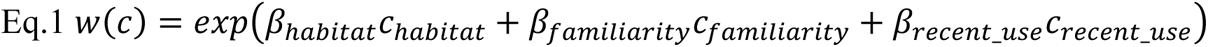

with w(c) the RSF score for the cell *c*. Habitat, familiarity and recent use are variables – described below – affecting the movement of the animals, with predetermined coefficients of selection that were varied, between simulations, from negative (i.e. simulating avoidance/repulsion), to positive (i.e. simulating selection/attraction) values. For all variables we used a gradient of values ranging from -4 to 4 by 0.5 increments.

##### ‘Habitat’ map

We assumed that two habitat types existed in the landscape. To create the habitat raster map, and for each replication of the landscape, we did as follows: using the localGibbs package (Michelot *et al*. 2019), we first generated over the extent of the map a spatially correlated Gaussian random field (with a defined spatial autocorrelation ρ=10) whose values, ranging between 0 and 1, where then attributed to each cell. We then used a threshold value *h* to discretize these continuous values into binary ones, so that 1 (respectively 0) represented the presence of habitat A (respectively B) in the cell. Thus, *h* also defined the proportion of habitat A in the landscape. By default, we used h=0.5. We initially tested the influence of the patchiness of the landscape by using two different values of spatial correlation when generating the Gaussian random field (see Figure S1 in Appendix S1 for illustration). Results were similar, and we therefore present here results obtained for the patchier landscapes.

##### ‘Familiarity’ map

Following previous work (Van Moorter *et al*. 2013; Wolf *et al*. 2009), we modelled familiarity as a binary raster map, with 0 (respectively 1) indicating an unfamiliar (respectively familiar) cell. During simulations, each individual had its own familiarity map, which was initialized at 0 for all cells. Thus, at the beginning of each simulation, individuals were assumed to be unfamiliar with the landscape. Then, at each time-step, the familiarity maps of all individuals were updated, with visited cells shifting to a value of 1. For simplicity, we did not include oblivion effect.

##### ‘Recent use’ map

During each simulation, the level of recent use of cells was tracked through one dynamic raster map, which was shared by all individuals of a simulation. At the beginning of the simulation, the map of recently used areas had cell values that were all 0. Then, at each time-step, once all individuals had relocated, we updated the map of recently used areas by calculating, for each cell, the number of times the cell had been visited by at least one individual over the last 20 time steps. This map can therefore be seen as storing an index of the recent presence of conspecifics. We used a temporal window since (1) animals may be able to detect only the recent, not the older, use of a location by other animals, such as from mark scents, and (2) since resources may regenerate after a long enough time period.

#### 2.1.2. Estimation of habitat selection coefficients from simulated movements

We used the model presented above to investigate to what extent selection for familiar or recently used locations could influence, and in particular possibly bias, the estimation of the strength of habitat selection when they are not taken into account.

We did so by, firstly, running 20 replications of the model for each combination of values of the selection coefficient for habitat, familiarity and recent use predictors. Secondly, we fitted, to each movement trajectory (i.e., 20 replications × 500 individuals), a RSF with habitat as the sole predictor, to obtain the estimate coefficient of selection for habitat 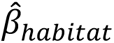 when familiarity and recently used areas were not known or taken into account. To do so, we calculated the selection ratio of each habitat 𝑆𝑅_𝐴_ (respectively 𝑆𝑅_𝐵_) as the proportion of cells of habitat A (respectively habitat B) among the used locations, divided by the proportion of cells of habitat A (respectively habitat B) among all cells of the landscape, i.e. the available locations. The habitat selection coefficient that would be estimated from a RSF with habitat as the sole predictor is given by (Eq. 2).

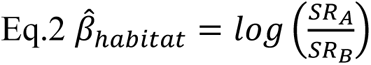

Lastly, we compared the estimated selection coefficient from the RSF 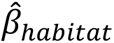 with the theoretical values in the simulation (𝛽_ℎ𝑎𝑏𝑖𝑡𝑎𝑡_), and investigated whether the difference between the two was important and varied consistently with the values of 𝛽_𝑓𝑎𝑚𝑖𝑙𝑖𝑎𝑟𝑖𝑡𝑦_ and 𝛽_𝑟𝑒𝑐𝑒𝑛𝑡𝑢𝑠𝑒_.

As we expected consequences on predictions of animal distribution (see section 2.2), we wanted to explore through which mechanism space familiarity and recent use of areas could impact these distributions. We focused on the movement rate of individuals, calculated here as the net displacement over 10 time steps, from simulations conducted in section 2.1.1. We used the average movement rates of individuals moving independently of familiarity and recent use (but with the same strength of habitat selection) as a reference value. We then compared, for all combination of values for the selection coefficients for familiarity and recent use predictors, the ratio between the average movement rates observed and the reference value.

### 2.2. Testing the effect of familiarity and recent use on population distribution predictions

As we found that not accounting for familiarity and recent use bias the estimation of habitat selection (see section 3.1), we studied to what extent this bias could lead to errors in the estimation of the distribution of the individuals in a changing environment.

First, to simulate a changing environment, we combined spatially correlated Gaussian random fields with planar gradient neutral landscapes to generate replicated series of four habitat maps that had an increasing proportion of the selected habitat along the West-East axis, which could therefore be seen as an axis of good habitat expansion. Each series was assumed to represent one changing landscape at four arbitrary times referred as T0, T1, T2, T3 (see Figure S1 in Appendix S2 for an illustration of one series).

Second, for each combination of coefficients for habitat type (𝛽_ℎ𝑎𝑏𝑖𝑡𝑎𝑡_), familiarity (𝛽_𝑓𝑎𝑚𝑖𝑙𝑖𝑎𝑟𝑖𝑡𝑦_) and recent use (𝛽_𝑟𝑒𝑐𝑒𝑛𝑡𝑢𝑠𝑒_), we ran simulations on landscape series, simulating the movement of 500 individuals over 500 time-steps in each landscape (T0 to T3). At the start of the simulations (at T0), individuals were randomly located within the western area (position on the x axis ≤ 10) where habitat A was more common. Then, their locations at the end of T0 were used at the beginning of T1 simulations, and so on. Simulations followed 4 scenarios differing by how animal movement was simulated:

(reference scenario) the movement of animals was based on a fully parametrized LG movement model whose coefficients were those used in section 2.1 (𝛽_ℎ𝑎𝑏𝑖𝑡𝑎𝑡_, 𝛽_𝑓𝑎𝑚𝑖𝑙𝑖𝑎𝑟𝑖𝑡𝑦_, 𝛽_𝑑𝑒𝑝𝑙𝑒𝑡𝑖𝑜𝑛_). This scenario was assumed to represent ‘the truth’.

(scenario 1) the movement of animals was influenced only by habitat type, based on a LG movement model whose coefficient for habitat type (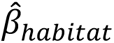) had been estimated in section 2.1 by fitting a habitat-only RSF model to movement data obtained by simulations with a fully parametrized LG model. Distributions emerging from this scenario are those that ecologists would often generate as they emerge from the standard approach of making predictions from estimations of habitat selection coefficients without accounting for ‘invisible landscapes’.

(scenario 2) the movement of animals was based on a fully parametrized LG model, as in reference scenario, but with the coefficient for habitat type being the one estimated from an habitat-only RSF model, as in scenario 1 (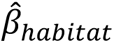𝛽_𝑓𝑎𝑚𝑖𝑙𝑖𝑎𝑟𝑖𝑡𝑦_𝛽_𝑟𝑒𝑐𝑒𝑛𝑡𝑢𝑠𝑒_). This scenario allowed us to test to what extent including correct information on the effect of ‘invisible landscapes’ allowed us to correct prediction errors induced by a biased estimation of selection for habitat type.

(scenario 3) the movement of animals was based on a habitat-only LG model, with the coefficient for habitat type being the correct one (𝛽_ℎ𝑎𝑏𝑖𝑡𝑎𝑡_). This scenario allowed us to test to what extent overlooking ‘invisible landscapes’ can induce errors in predictions, if selection for habitat type has been correctly estimated.

For all scenarios, simulations were replicated 20 times, and for all simulations, we estimated at time T0, T1, T2, T3 the density of animal locations across the West-East axis of good habitat expansion. This allowed us to describe the changes in the distribution of the population in the landscapes. To compare the distributions emerging from scenario 1 to 3 with the ones from the reference scenario, we used the index proposed by (Pastore & Calcagnì 2019) to measure the overlap between the distributions. The index takes greater values as overlap increases, up to a maximum value of 1 when the compared distributions are exactly the same.

## 3. RESULTS

### 3.1. Influence of ‘invisible landscapes’ on habitat selection analysis

Estimating the strength of habitat selection (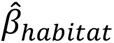) from a habitat-only model clearly provided biased results when the animal movement was driven by habitat selection but also other variables such as familiarity (Fig. 1a), or recent use (Fig. 1b). Importantly, our results show that habitat selection is estimated to be stronger than it actually is if the animal tends to select for greater value of the overlooked variables, i.e. if the selection coefficients for these variables are positive. The opposite hold true for negative coefficients. The bias increases when the influence of these overlooked variables on movement increases (i.e. when coefficients are greater). We observed that the biases associated with each overlooked variable could sometimes compensate each other, as habitat selection strength was correctly estimated for specific combinations of non-null 𝛽_𝑓𝑎𝑚𝑖𝑙𝑖𝑎𝑟𝑖𝑡𝑦_ and 𝛽_𝑟𝑒𝑐𝑒𝑛𝑡𝑢𝑠𝑒_ (see Figure S2 in Appendix S1). Note that, when the animal’s movement was not influenced by familiarity or recent use, we could correctly recover, with some noise due to the stochastic nature of simulations, the theoretical coefficient of habitat selection used in the model. This is expected but demonstrates the validity of the approach. In addition to the bias observed in the estimation of the strength of selection for habitat variables, we found that individuals selecting familiar habitats exhibited smaller movement rates than individuals moving independently of familiarity (Fig. 2a). In contrast, we did not find any influence of the selection, nor the avoidance, of recently used habitats on the movement rate of the simulated individuals (Fig. 2b).

**Figure 1.**
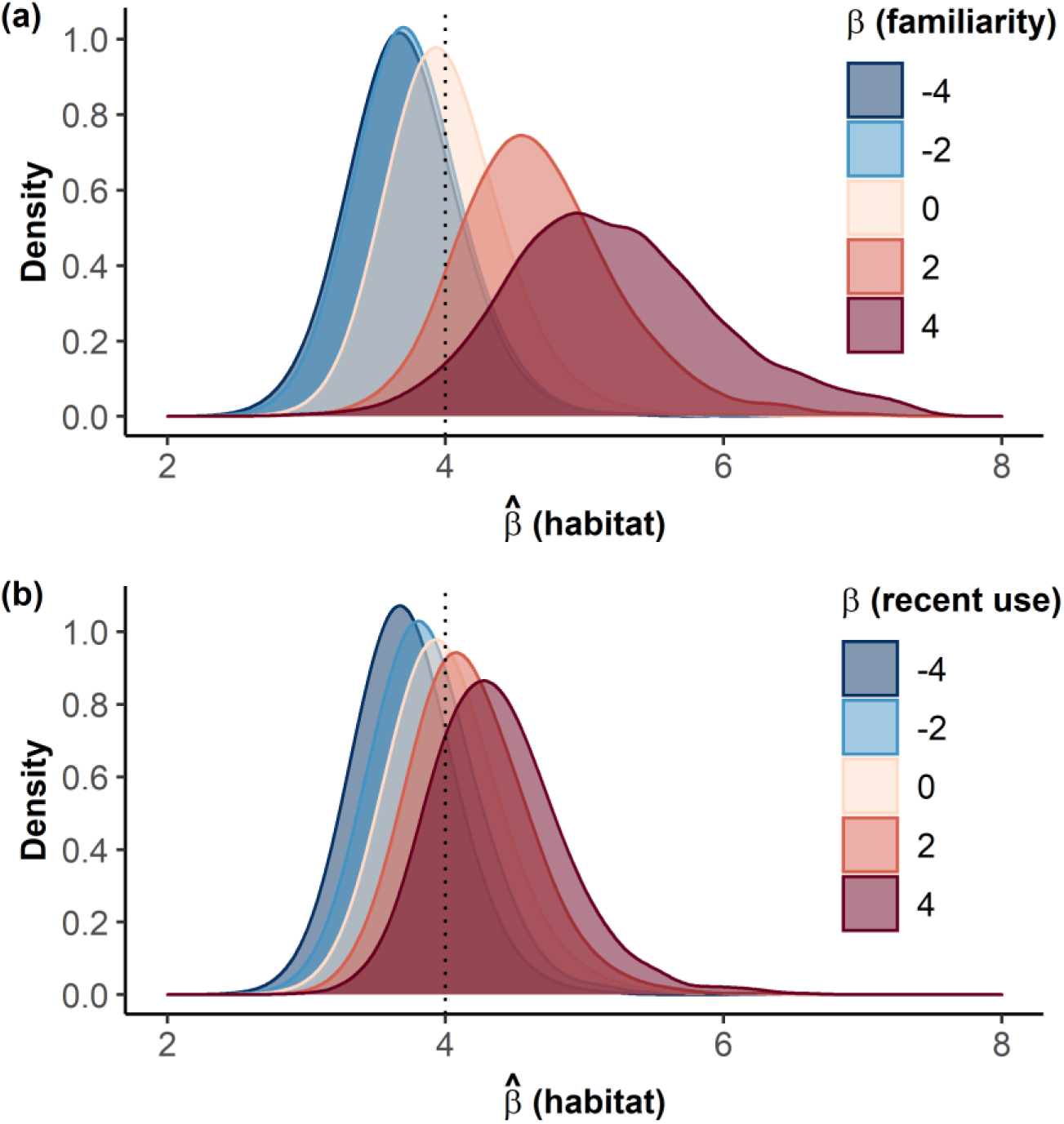
Influence of the strength of selection for familiar space (𝛽_𝑓𝑎𝑚𝑖𝑙𝑖𝑎𝑟𝑖𝑡𝑦_; A) and for recently used areas (𝛽_𝑟𝑒𝑐𝑒𝑛𝑡𝑢𝑠𝑒_; B) on the estimation of the strength of selection for habitat type (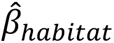). The dotted vertical lines indicate the value of 𝛽_ℎ𝑎𝑏𝑖𝑡𝑎𝑡_ used in the simulations. The distribution of the estimates 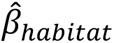 across varying values of 𝛽_𝑓𝑎𝑚𝑖𝑙𝑖𝑎𝑟𝑖𝑡𝑦_ and 𝛽_𝑟𝑒𝑐𝑒𝑛𝑡𝑢𝑠𝑒_ are shown along a blue (i.e. negative strength) to red (i.e. positive strength) gradients. The influence of space familiarity on habitat selection estimations is shown for null effect of recent use variables (𝛽_𝑟𝑒𝑐𝑒𝑛𝑡𝑢𝑠𝑒_ = 0; A), and inversely (𝛽_𝑓𝑎𝑚𝑖𝑙𝑖𝑎𝑟𝑖𝑡𝑦_ = 0; B). Simultaneous influences of familiarity and recent use variables are provided in Figure S2 in Appendix S1.

**Figure 2.**
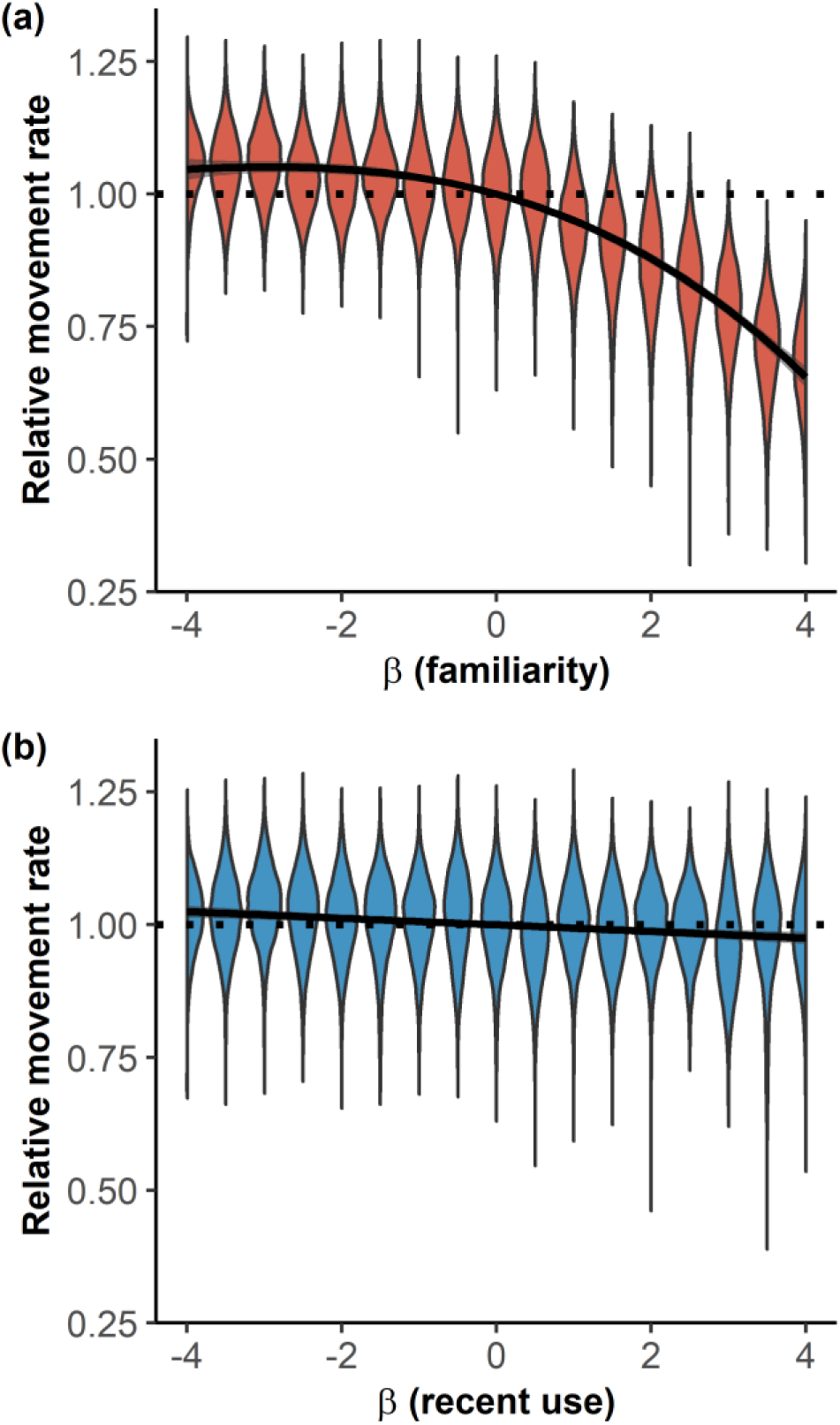
Influence of the strength of selection for familiar space (𝛽_𝑓𝑎𝑚𝑖𝑙𝑖𝑎𝑟𝑖𝑡𝑦_; A) and for recently-used areas (𝛽_𝑟𝑒𝑐𝑒𝑛𝑡𝑢𝑠𝑒_; B) on the movement rate of simulated individuals, relatively to the average movement rate of individuals with same selection strength for habitat type, but with null selection for familiar space and recently use areas. Violin plots show the distribution of movement rates obtained over 20 habitats × 500 replications for each set of habitat selection coefficients for familiar and recent use habitats.

### 3.2. Influence of ‘invisible landscapes’ on population-level inference

Predictions of the distribution of individuals in a changing landscape, when made using models previously fitted with only habitat as predictor, were increasingly wrong over time (Fig. 3; Fig. 4; scenario 1). This was particularly true when the selection of animal for familiar areas was important, and to a lesser extent when the selection for areas recently used by conspecifics was also important (Fig. 4). In any case, the error made was a consistent bias towards an over-estimation of the tracking of the change in the distribution of the good habitat by the population when individuals selected for familiar locations (Fig. 3). Conversely, we would under-estimate the tracking of the change of the distribution of the good habitat by the population when individuals avoid familiar or recently use locations (Figure S2 in Appendix S2).

**Figure 3.**
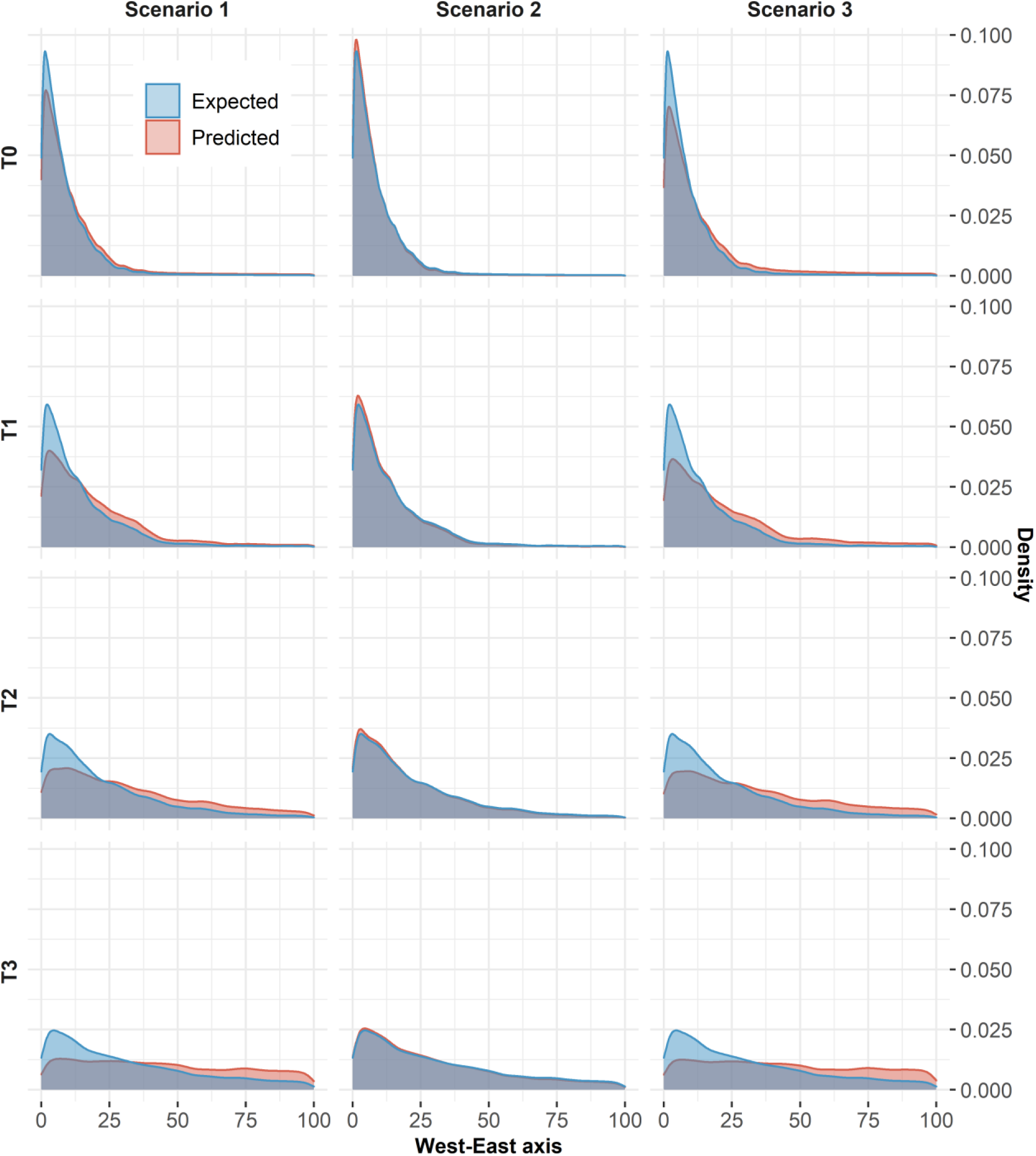
Comparison between the expected (reference scenario; blue) and the predicted (scenario 1; 2; 3; red) distributions of individuals along a habitat-expanding gradient (West-East axis) at time T0, T1, T2, T3. The expected distribution was obtained by simulating individuals with a fully parametrized LG movement model (𝛽_ℎ𝑎𝑏𝑖𝑡𝑎𝑡_ = 4; 𝛽_𝑓𝑎𝑚𝑖𝑙𝑖𝑎𝑟𝑖𝑡𝑦_ = 2; 𝛽_𝑟𝑒𝑐𝑒𝑛𝑡𝑢𝑠𝑒_ = 0). Scenario 1: distribution predicted from the biased estimate of habitat selection (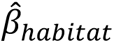 = 4.7; 𝛽_𝑓𝑎𝑚𝑖𝑙𝑖𝑎𝑟𝑖𝑡𝑦_ = 0; 𝛽_𝑟𝑒𝑐𝑒𝑛𝑡𝑢𝑠𝑒_ = 0). Scenario 2: distribution predicted from biased estimate of habitat selection and the selection coefficients for familiar and recently used areas (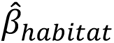 = 4.7; 𝛽_𝑓𝑎𝑚𝑖𝑙𝑖𝑎𝑟𝑖𝑡𝑦_ = 2; 𝛽_𝑟𝑒𝑐𝑒𝑛𝑡𝑢𝑠𝑒_ = 0). Scenario 3: distribution predicted from the coefficient of habitat selection only (𝛽_ℎ𝑎𝑏𝑖𝑡𝑎𝑡_ = 4; 𝛽_𝑓𝑎𝑚𝑖𝑙𝑖𝑎𝑟𝑖𝑡𝑦_ = 0; 𝛽_𝑟𝑒𝑐𝑒𝑛𝑡𝑢𝑠𝑒_ = 0).

**Figure 4.**
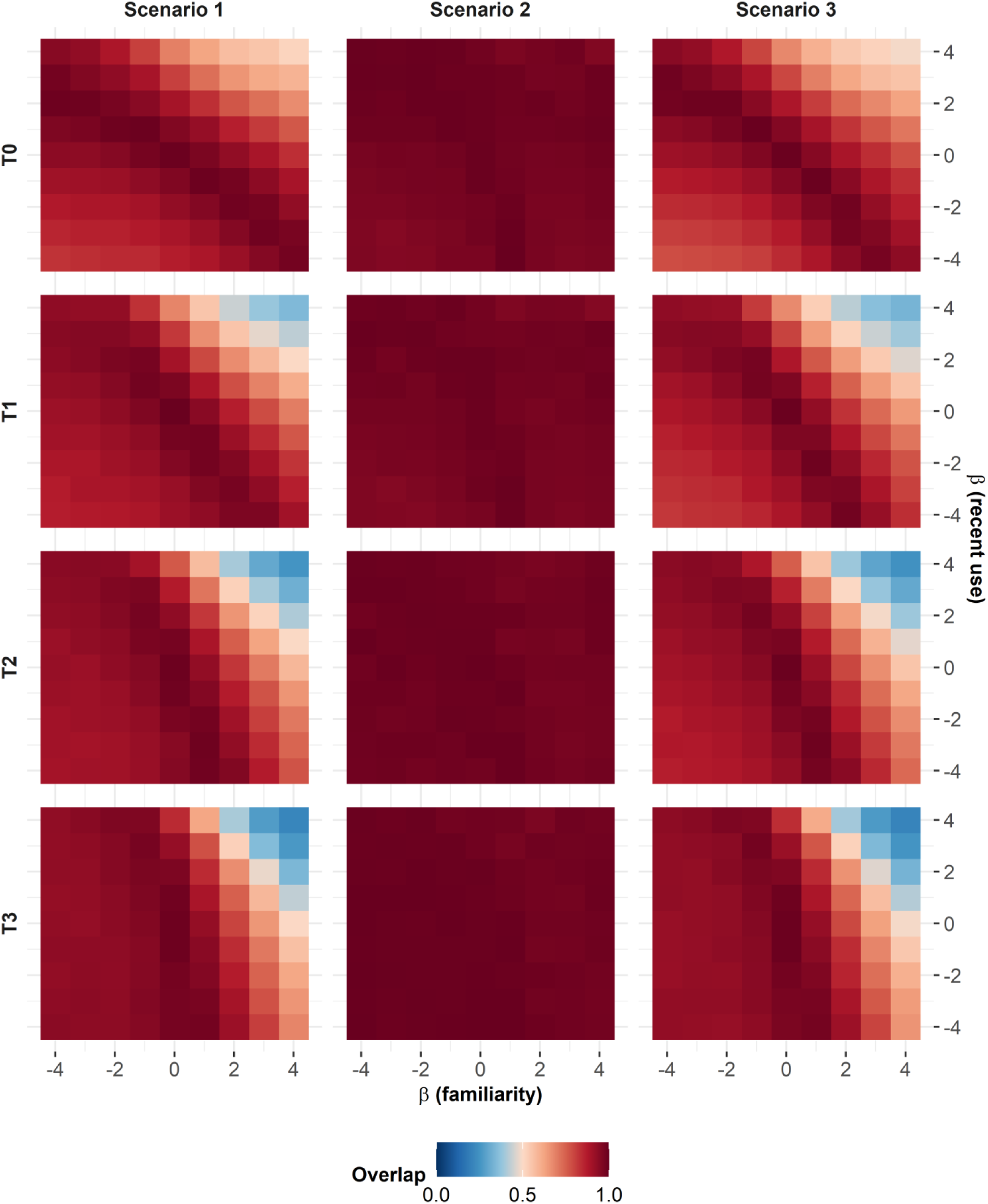
Overlap between the predicted and expected population distributions along a habitat-expanding gradient (T0-T4) according to the individual selection coefficient for familiar space (𝛽_𝑓𝑎𝑚𝑖𝑙𝑖𝑎𝑟𝑖𝑡𝑦_) and recently used areas (𝛽_𝑟𝑒𝑐𝑒𝑛𝑡𝑢𝑠𝑒_) used in the expected distribution case. Scenario 1: Population distributions predicted from the estimated coefficient of habitat selection (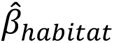). Scenario 2: Population distributions predicted from the estimated coefficient of habitat selection and the selection coefficients for familiar and recently used habitats (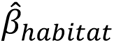, 𝛽_𝑓𝑎𝑚𝑖𝑙𝑖𝑎𝑟𝑖𝑡𝑦_, 𝛽_𝑟𝑒𝑐𝑒𝑛𝑡𝑢𝑠𝑒_). Scenario 3: Population distributions predicted from the actual coefficient of habitat selection (𝛽_ℎ𝑎𝑏𝑖𝑡𝑎𝑡_)

Somewhat unexpectedly, predictions made with models that had the correct selection coefficients for familiarity and recent use variables, but for which the coefficient for the habitat type variable had been estimated from habitat-only models as in scenario 1, were most often good (Fig. 4; scenario 2). Predictions made with models that had the correct selection coefficient for habitat type but did not include familiarity or recent use variables were often wrong (Fig. 4; scenario 3), in a way similar to what was observed in scenario 1.

## 4. DISCUSSION

In this study, we used computer simulations to show that overlooking ‘invisible landscapes’, i.e., non-physical variables that are relevant for animals (competition, social attraction, space familiarity) but difficult to grasp and measure for the human analyst, induces bias in habitat selection analyses and population-level projections. These results are important because an increasing number of empirical studies are confirming what had long been ‘known’ by field ecologists, that such processes (e.g. attraction to familiar sites or to social mates) are indeed important, if not critical, in driving animals’ movement decisions (Merkle *et al*. 2014, 2019; Peignier *et al*. 2019; Ranc *et al*. 2020; Wolf *et al*. 2009).

### 4.1. Why do some landscapes remain ‘invisible’ to human analysts?

Of the ‘invisible landscapes’ that affect the habitat choices of animals, not all present the same challenges to the human analyst. Numerous methods now allow to gain information about the current and past presence of social mates, competitors and predators: for instance, remote observations using camera-traps or passive acoustic loggers can provide relevant data that can be used to create proxy variables, whose quality will likely depend on the monitoring effort deployed (e.g. density of recorders) (Caravaggi *et al*. 2017; Gibb *et al*. 2019). In some rare instances, enough competitors and predators can be tracked by GPS to provide similar information (Wilmers *et al*. 2015). Consideration of such variables in models based on tracking data has fortunately been increasing in the last decades, for example by adding a covariate to describe the familiarity of each used/available locations in habitat selection analysis (Kim *et al*. 2023; Oliveira-Santos *et al*. 2016) or a covariate indexing the likelihood of presence of predators (Davies *et al*. 2021; Willems & Hill 2009). Unfortunately, animals’ decisions *in natura* are likely driven by a number of these non-physical variables, often turning them as ‘invisible landscapes’ because of their difficulty to grasp and measure. Considering larger areas to account for a greater diversity of habitats may come at the expense of being less able to accurately account for some of the ‘invisible landscapes’ that may be experienced by animals at finer spatial scales. One key remark we wish to make here is that information about a number of important factors driving animal’s decisions about where to move, such as space familiarity, past experience with predators, or more generally any variable related to the past experience of a specific individual, will never be available in studies only based on snapshot (one-point-in-time) observations of species’ presence. Our results provide general guidance on how systematic biases are therefore expected from habitat selection analyses (see section 4.2), and by consequence species distribution models developed from such species observation data (see section 4.3).

### 4.2. What are the consequences of overlooking ‘invisible landscapes’ when estimating habitat selection?

Overlooking the attraction of animals towards ‘invisible landscapes’, for instance familiar space, consistently led to an over-estimation of the strength of habitat selection. This result arose from a positive feedback loop that makes animals over-using good habitats: by being more familiar with good habitats (because of a preliminary habitat selection), animals that randomly select a location among the set of familiar locations are more likely to draw a location from good habitats. Similar results – and feedback loops – were obtained regarding the ‘recent use’ variable: overlooking the avoidance towards recently used areas led to under-estimations of animal’s habitat selection. Interestingly, the reverse mechanism had been described by (Van Moorter *et al*. 2013), showing how one can estimate a spurious selection for familiar places what is the selection for a physical variable lacking from the habitat selection analysis. Together, these results show the importance of exhaustively describing the various ‘landscapes’ on which an animal bases its decision to move when estimating habitat selection. Therefore, results from SDMs based on snapshot observation of unknown individuals that have no other choice than to overlook the effect of attraction for familiar places – a likely very common process (Merkle *et al*. 2014, 2019; Ranc *et al*. 2020; Wolf *et al*. 2009) – should be taken with caution, and should be valid only if the influence of physical variables, relatively easily mapped, is largely dominant over the one of the ‘invisible landscapes’. This remains to be properly evaluated by behavioural ecologists and will likely be best done by comprehensive studies based on tracking individuals while monitoring as closely as possible their environment, including conspecifics, competitors, and predators.

### 4.3. How to integrate ‘invisible landscapes’ in species distribution models?

Accurately estimating habitat selection is obviously an important step toward unbiased projections of population distribution. However, here, we found that mechanistic distribution modelling integrating the cognitive processes underlying animal movement is, at least, as important as accurately estimating habitat selection. Indeed, our results show that integrating ‘invisible landscapes’ not only when building SDM/RSF explanatory models to obtain coefficients, but also when using these models for predictions is crucial to accurately predict population distribution, and can even mitigate the consequence of estimating biased coefficients of habitat selection (see section 3.2). As a result, if SDMs based on snapshot observation of unknown individuals fail to integrate ‘invisible landscapes’ when linking species occurrence to landscape composition, efforts should be done to integrate ‘invisible landscapes’ in the projections made from these models. In particular, our results suggest that the reduction of the movement rate of animals using spatial memory to navigate is an important factor to consider when predicting the distribution of animals over changing conditions. Unfortunately, there does not seem to be any obvious solution to this important issue. The SDM literature has recently seen a surge of methods to account for dispersal and connectivity constraints (Barbet-Massin *et al*. 2012; Boulangeat *et al*. 2012; Monsimet *et al*. 2020), that could provide a partial solution to this problem, although ultimately those does not represent directly the process presented here.

Integrating past experience will likely require using mechanistic SDM that directly simulate biological processes (as opposed to correlative SDM; sensu Dormann et al. 2018). Thus, we add to the list of reasons (e.g. integration of physiological processes or biotic interaction) that have been called for to move away from correlative SDM. While doing so, we recognize the complexity and challenges of parametrizing mechanistic models, but highlight that habitat selection models such as RSF or movement models such as integrated step-selection functions or LG models are appropriate tools to do gain insights about parameters. Ultimately, predictive distribution models should also include the demographic fate of individuals, and the recently developed spatial absorbing Markov chain framework (Fletcher *et al*. 2019) is an appealing first step towards this goal, of intermediate complexity. A recent application even accounts for site fidelity (Ditmer *et al*. 2023), although fidelity remains linked to habitat quality rather than past experience of the individuals.

## 5. CONCLUSION

The estimation of the strength of habitat selection for specific, easily measurable variables, such as vegetation cover types, is at the heart of the species distribution models on which we, ecologists, base our prediction of animal distribution under changing environments. Yet, we have shown here that bias in the estimation easily arise when one does not, or cannot, account for some ‘invisible’ variables that affect the movement of animals, in particular familiarity with the place. We therefore urge researchers to go beyond the consideration of physical variables as predictors of movements (e.g. vegetation cover type, distance to features). Certainly, ecologists should strive to find ways to account for the effects of ‘invisible landscapes’, even if in imperfect ways. The importance of past experience of individuals, which is inaccessible to the human analyst using snapshot observation data of unknown individuals, calls for mechanistic distribution modelling integrating the cognitive processes underlying animal movement.

## Acknowledgements

We thank Marion Valeix who provided constructive comments on a previous draft.

## Statement of authorship

SCJ came up with the original idea. RLP and RD provided further critical insights to develop the conceptual framework. RD built the model and performed simulations. All authors interpreted the results. RD wrote the initial draft, to which all authors contributed substantial changes until submission.

## Appendix S1. Testing the influence of invisible landscapes on habitat selection analysis

**Figure S1.**
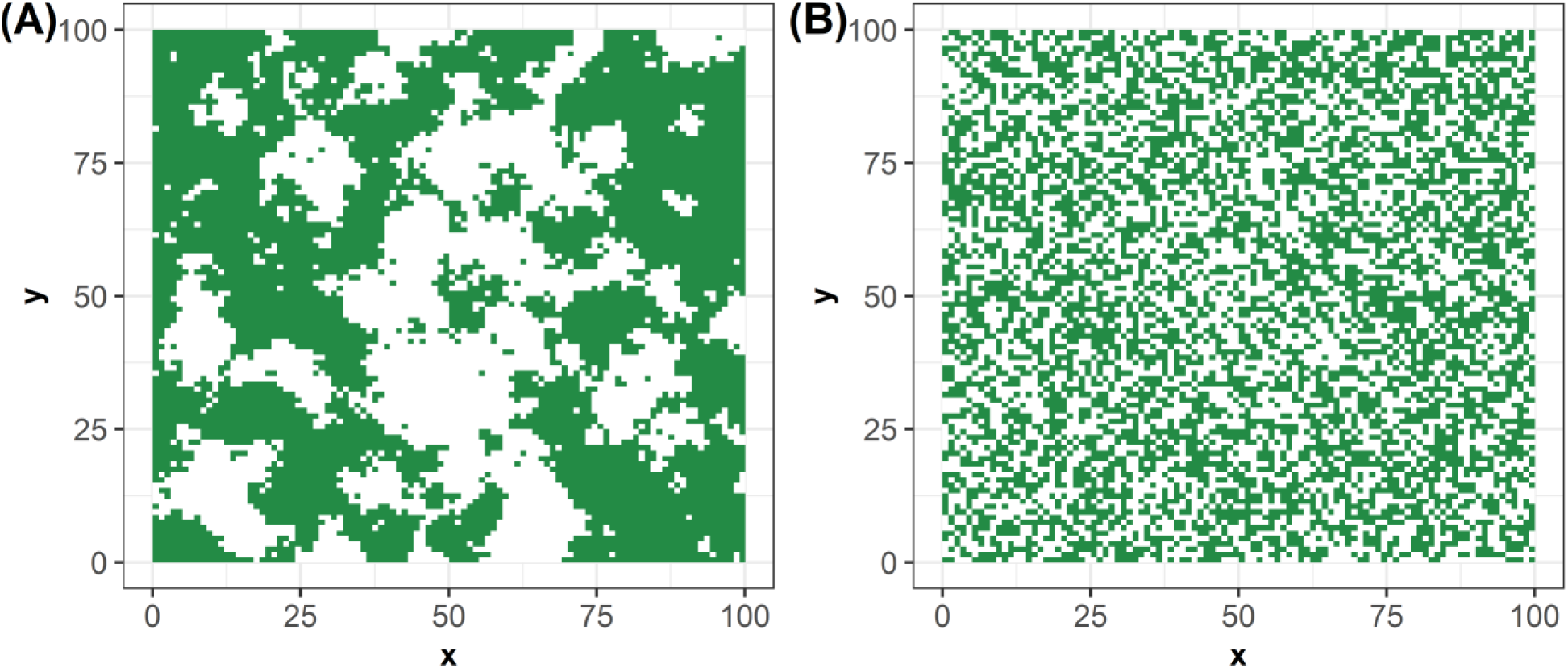
Example of landscapes simulated by discretizing spatially correlated Gaussian random fields. Green cells show the presence of the selected habitat-type (habitat A in main text). (A) More-patchy (ρ=10) and (B) less-patchy landscapes (ρ=1) were generated to test the influence of invisible landscape on habitat selection analysis.

**Figure S2.**
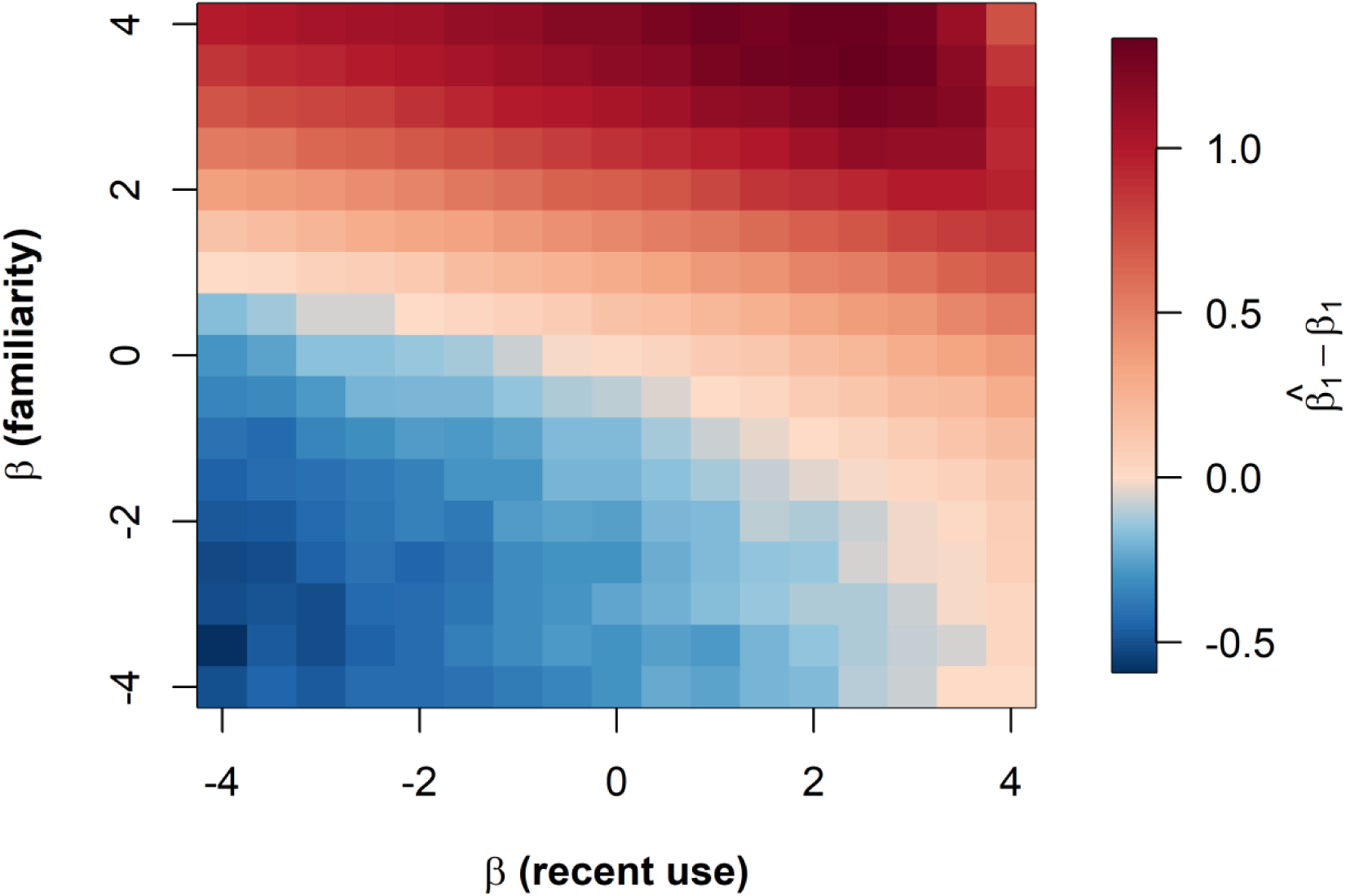
Influence of the strength of selection for familiar space (𝛽_𝑓𝑎𝑚𝑖𝑙𝑖𝑎𝑟𝑖𝑡𝑦_) and for recently-used areas (𝛽_𝑟𝑒𝑐𝑒𝑛𝑡 𝑢𝑠𝑒_) on habitat selection analyses. Cell values show the differences between the estimated (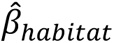) and expected 𝛽_ℎ𝑎𝑏𝑖𝑡𝑎𝑡_ coefficient of selection for habitat-type. Movement trajectories (n = 20 replications × 500 individuals ; step = 1000) were simulated on a fully parametrized LG movement model with coefficients (𝛽_ℎ𝑎𝑏𝑖𝑡𝑎𝑡_ = 4; 𝛽_𝑓𝑎𝑚𝑖𝑙𝑖𝑎𝑟𝑖𝑡𝑦_; 𝛽_𝑟𝑒𝑐𝑒𝑛𝑡 𝑢𝑠𝑒_).

## Appendix S2. Testing the influence of invisible landscapes on population distribution predictions

**Figure S1.**
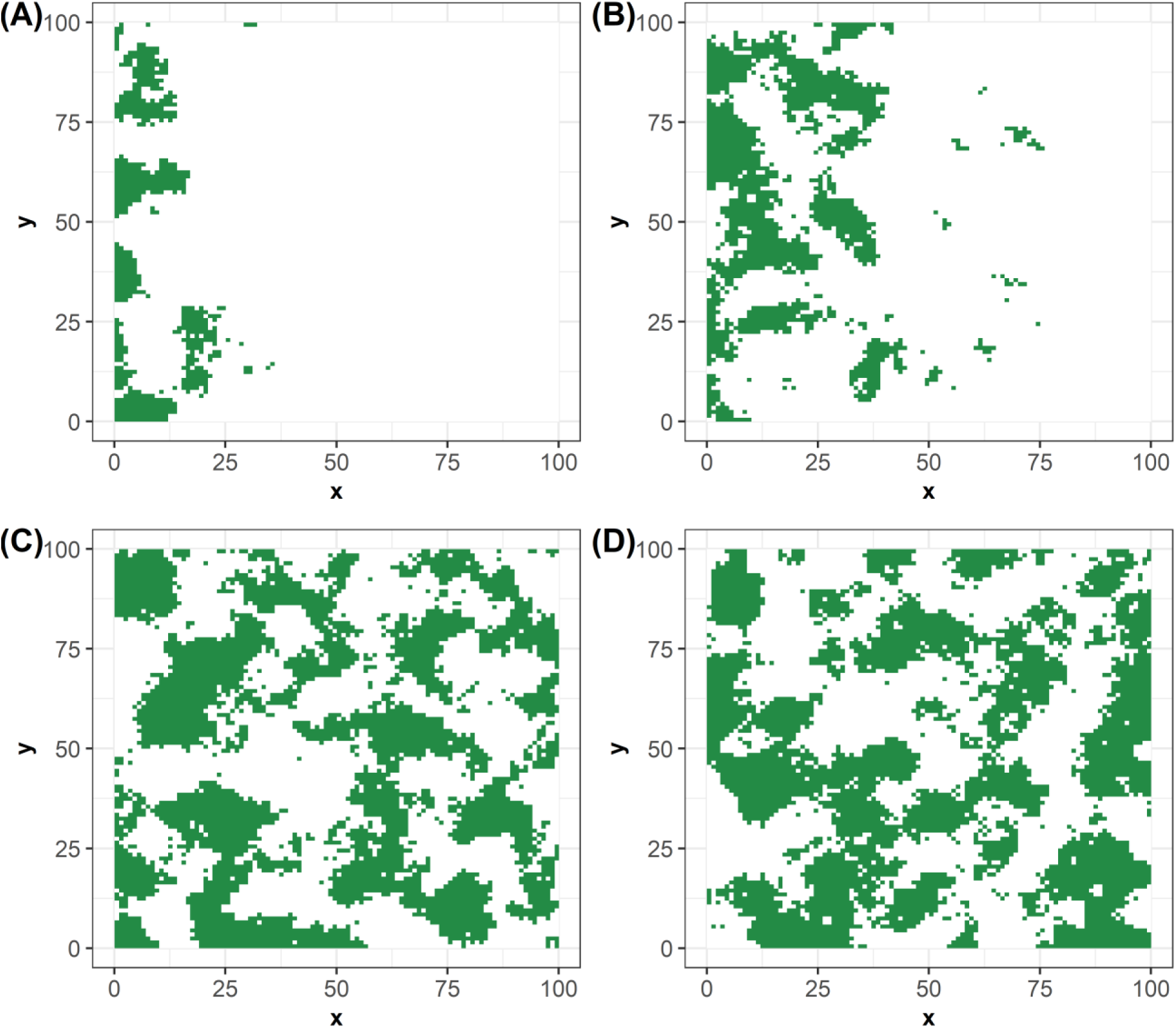
Example of landscapes simulated by discretizing spatially correlated Gaussian random fields, combined with planar gradient neutral landscapes with increasing proportions of the selected habitat (habitat A, in green), to generate four successive landscapes over time, referred as (A) T0, (B) T1, (C) T2, (D) T3 in the result section.

**Figure S2.**
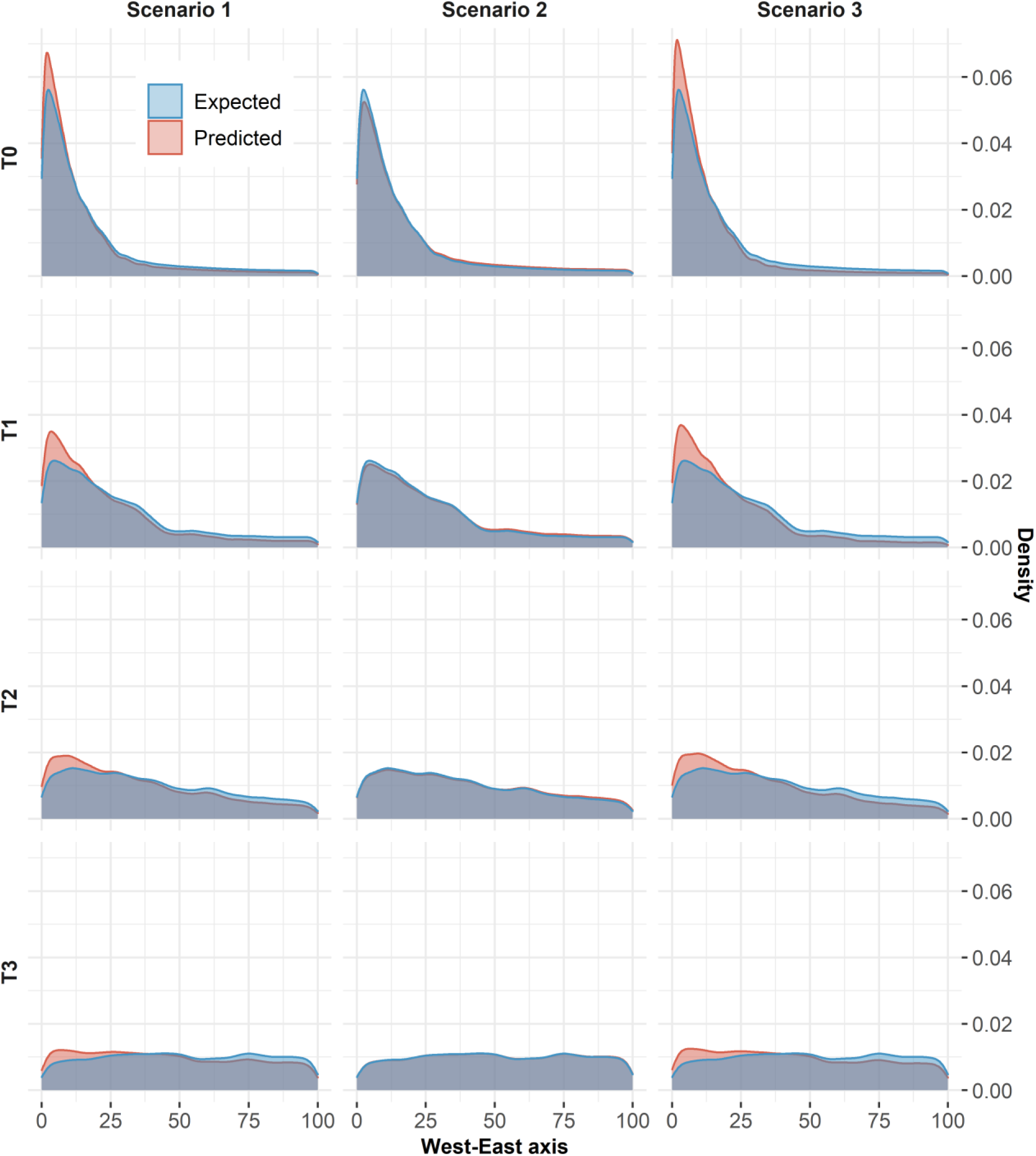
Population-distribution expected and predicted when simulated movements were independent on habitat familiarity (𝛽_𝑓𝑎𝑚𝑖𝑙𝑖𝑎𝑟𝑖𝑡𝑦_ = −4) and habitat depletion (𝛽_𝑑𝑒𝑝𝑙𝑒𝑡𝑖𝑜𝑛_ = 0).

